# Artificial Intelligence based Liver Portal Tract Region Identification and Quantification with Transplant Biopsy Whole-Slide Images

**DOI:** 10.1101/2022.08.31.506101

**Authors:** Hanyi Yu, Nima Sharifai, Kun Jiang, Fusheng Wang, George Teodoro, Alton B. Farris, Jun Kong

## Abstract

Liver fibrosis staging is clinically important for liver disease progression prediction. As the portal tract fibrotic quantity and size in a liver biopsy correlate with the fibrosis stage, an accurate analysis of portal tract regions is clinically critical. Manual annotations of portal tract regions, however, are time-consuming and subject to large inter- and intra-observer variability. To address such a challenge, we develop a Multiple Up-sampling and Spatial Attention guided UNet model (MUSA-UNet) to segment liver portal tract regions in whole-slide images of liver tissue slides. To enhance the segmentation performance, we propose to use depth-wise separable convolution, the spatial attention mechanism, the residual connection, and multiple up-sampling paths in the developed model. This study includes 53 histopathology whole slide images from patients who received liver transplantation. In total, 6,012 patches derived from 30 images are used for our deep learning model training and validation. The remaining 23 whole slide images are utilized for the model testing. The average liver portal tract segmentation performance of the developed MUSA-UNet is 0.94 (Precision), 0.85 (Recall), 0.89 (F1 Score), 0.89 (Accuracy), 0.80 (Jaccard Index), and 0.91 (Fowlkes–Mallows Index), respectively. The clinical Scheuer fibrosis stage presents a strong correlation with the resulting average portal tract fibrotic area (R=0.681, p<0.001) and portal tract percentage (R=0.335, p=0.02) computed from the MUSA-UNet segmentation results. In conclusion, our developed deep learning model MUSA-UNet can accurately segment portal tract regions from whole-slide images of liver tissue biopsies, presenting its promising potential to assist liver disease diagnosis in a computational manner.

## 1. INTRODUCTION

Detection of early stage fibrosis in transplant liver biopsies is important for predicting disease progression and guiding medical management^1^. Known as a strong predictor of liver disease progression and mortality, liver fibrosis can be captured by multiple non-invasive medical imaging techniques, such as computed tomography (CT), magnetic resonance elastography (MRE), and transient elastography (TE)^2^. For accurate liver fibrosis staging, however, the histopathologic examination of liver biopsy samples remains the “gold standard” for liver fibrosis assessment^1^. Although numerous histopathological staging systems have been utilized for liver fibrosis evaluation in current clinical practice, including Knodell, Metavir, Ishak, and Scheuer systems, only manual reviews or semi-quantitative evaluations are conducted by these staging systems, resulting in large inter- and intra-observer variability^3-5^.

To reduce such variations, evaluation methods based on artificial intelligence (AI) algorithms, such as random forests, K-nearest neighbors, and support vector machines, have been developed to provide objective diagnostic tools for liver fibrosis staging^6-8^. In contrast to these conventional machine learning methods, deep learning has emerged as a powerful tool for diverse biomedical image processing studies due to its great success across different image modalities^9^. The family of deep learning methods originated from artificial neural networks that consist of layers of computational nodes analogous to neurons in human brains^10,11^. Deep Convolutional Neural Networks (DCNNs) are a class of deep learning methods where convolution filters in different layers extract image features at different resolution levels^12^. Unlike the traditional machine learning methods, DCNNs require no manual feature engineering and can support multiple imaging modalities, including CT^13,14^, MRI^2^, and ultrasonography images^15,16^. The resulting image features and other clinical demographic information (e.g., gender and age) can be leveraged for an integrated prediction analysis by multiple fully connected layers attached to the DCNN backbone.

Deep neural networks are also powerful AI tools for the semantic segmentation analysis of a large spectrum of biomedical images. Although it used to be time-consuming for a DCNN model to produce a pixel by pixel semantic segmentation map^17^, Fully Convolutional Network (FCN)^18^ has been developed to improve the processing speed. As a significant breakthrough allowing for an efficient segmentation map generation, FCN has presented its high value for radiography, ultrasonography, and histopathological image segmentation^19^. As the deep learning techniques evolve, Mask-RCNN^20^ has been proposed to combine the FCN with the Faster R-CNN^21^ model. Among other applications^22,23^, Mask-RCNN has been successfully applied to recover clumped steatosis droplets in liver histopathological images^24^. Leveraging the FCN as a building block, the UNet architecture^25^ and its extensions, in turn, have become the widely used deep learning models for biomedical image segmentation analysis. Recently, two UNet models were cascaded to solve a segmentation task where the first UNet cropped volumes of interest (VOIs) from full-resolution 3D CT image volumes and the second UNet classified voxels within such VOIs^26^. Additionally, two UNet architectures sharing the same encoder were built for the prediction of two separate segmentation maps, one for cell nuclei and another for boundaries^27^. The resulting two segmentation maps were next synthesized for the final segmentation results. In another study on human skin, the convolution layer sequences in the original UNet architecture were replaced with dense-connected blocks to improve the segmentation performance with multiphoton microscopy images of in vivo human skins at the expense of doubled convolution layer number^28^. Furthermore, residual connections to convolution layer sequences were also proposed to outperform the original UNet with fluorescence microscopy, dermoscopy, endoscopy, and MR images^29^.

Inspired by the promising deep learning performance, some studies have been carried out for fibrosis analysis with liver whole-slide images (WSIs). Although a study used a pre-trained AlexNet^30^ to predict the liver fibrosis stage, its input images were acquired from second-harmonic generation microscopy^5^. A modified UNet architecture was also utilized to detect portal tract regions in mouse liver biopsy histopathology WSIs, but no comparison experimental result was given^31^. In our prior work^1^, we have manually delineated portal tract regions in liver biopsy images and demonstrated that the resulting quantitative portal tract fibrotic percentage and average portal tract area of portal tract regions are correlated with the liver fibrosis stage made by domain experts. However, such results are subject to intra- and inter-observer variability due to the manual annotation process^3^. Therefore, the development of fully automated and accurate segmentation algorithms for liver portal tract regions is an essential step to improve the evaluation consistency.

In this study, we propose a novel UNet-based deep convolutional neural network that automatically segments portal tract regions from high-resolution liver biopsy WSIs. To enhance performance, we substitute the two cascaded convolution structures in the original UNet design with a Residual Spatial Attention (RSA) processing block. Additionally, the output layer of our developed network directly synthesizes up-sampling features from multiple image resolutions. By such a Multiple Up-sampling Path (MUP) mechanism, the developed deep learning model reduces the false-negative rate and generates smoother borders. The network is trained with image patches and applied to liver biopsy WSIs. The resulting portal tract fibrotic percentage and average portal tract fibrotic area computed by our method present a strong correlation with the clinical Scheuer fibrosis stage. The performance of our approach is both qualitatively and quantitatively compared with that of the widely used methods. To demonstrate the contribution from individual modules, we also conduct ablation experiments and present ablation study results. Our method presents superior performance, suggesting its promising potential to assist clinical diagnosis.

## 2. MATERIALS AND METHODS

We present the overall schema of the proposed study in Figure 1(A). Images in our dataset are scanned with stained liver biopsy sections and utilized for training our proposed deep neural network. With network prediction results and human annotations, we quantitatively evaluate the network performance by statistical analyses. The Emory University Institutional Review Board approved these studies and waived the need for informed consent (IRB # CR002-IRB00055904).

**Figure 1.**
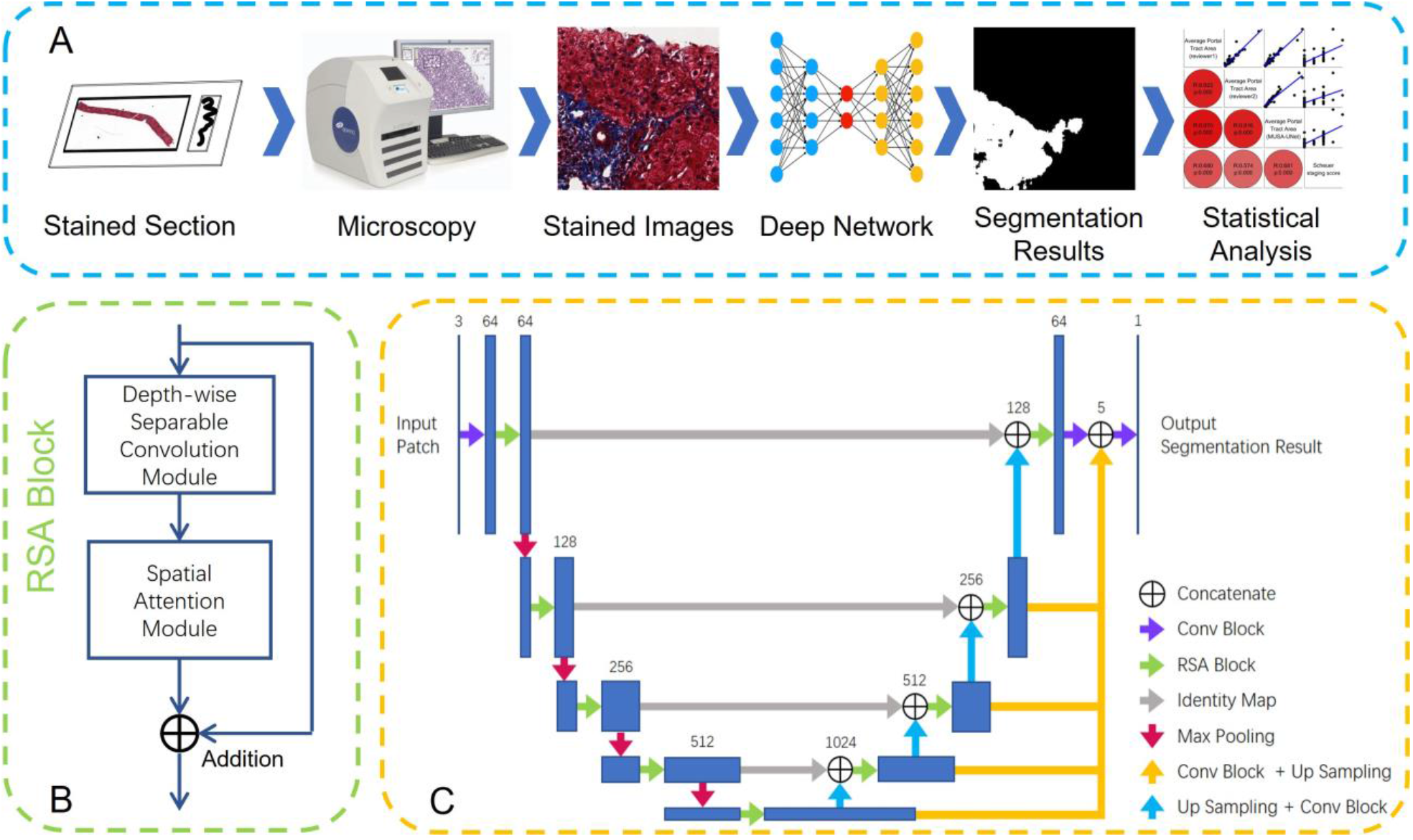
Overall schema of our proposed model. (A) Tissue sections were fixed, embedded, stained, and scanned for WSI generation. Resulting WSIs with human annotations are provided to our proposed MUSA-UNet for portal tract segmentation and statistical analyses; (B) We present the structure of the proposed RSA block that substitutes cascaded convolutional layers in the traditional UNet architecture. It primarily consists of one Depth-wise Separable Convolution (DSC) block and one Spatial Attention (SA) module connected by a residual network; (C) Our proposed deep learning neural network MUSA-UNet for image segmentation concatenates features from all decoders. As there are multiple paths providing lower resolution features from decoders to the output layer, such a Multiple Up-sampling Path (MUP) mechanism alleviates the false negative problem noticeable in the original UNet model in our study.

### 2.1 Tissue preparation

A retrospective study at Emory University was conducted on liver biopsies from patients who had undergone liver transplantation. Liver biopsy specimens were fixed in 10% buffered formalin and embedded in paraffin. Histologic sections were cut at 5 μm and stained with Masson’s trichrome and hematoxylin and eosin (H&E). All sections were staged histologically by pathologist visual assessments using the criteria of Scheuer score, ranging from 0 to 4. Biopsy specimens that pathologists could not stage due to either inadequate materials or documented fragmentations in the final pathological report were excluded. Biopsy specimens presenting the material adequacy are those that have a core length of at least 10 mm and at least 5 portal tracts. Trichrome-stained sections were scanned by an Aperio ScanScope CS (Aperio Technologies Inc., Vista, CA). The scanning was performed at 40x (i.e., 20x with 2x magnification doubler) with a numerical aperture of 0.75, giving a 40x resolution of 0.25 μm/pixel.

### 2.2 Deep neural network architecture

To make a full use of image information for segmentation, we have developed a Multiple Up-sampling and Spatial Attention guided UNet model (MUSA-UNet) that leverages the UNet architecture as the building block. The UNet architecture is known as a symmetric encoder-decoder framework that can effectively differentiate foreground pixels from the background by learning and incorporating local features from the higher resolution images and global information from the lower resolution images^25^. However, we notice UNet demonstrates a noticeably high false-negative rate by our experiments. To enhance model performance, we design two new mechanisms to specifically address this problem.

(1) We have developed a new Residual Spatial Attention (RSA) block to replace the sequence of two convolution layers in the original UNet for enhanced network performance. We propose the RSA block that consists of a residual network embedded with one Depth-wise Separable Convolution (DSC) and one Spatial Attention (SA) module. The RSA block architecture is presented in Figure 1(B). Specifically, the output of a RSA block can be formulated as follows:

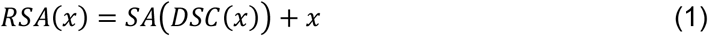

where *x* is the input feature array; *SA*(·)and *DSC*(·)are the spatial attention and the DSC module, respectively.

A DSC module has been proposed to divide a regular convolution layer into a depth-wise and a point-wise convolution layer for parameter number regulation^32,33^. It has been shown that the performance of a DSC module is similar to that of the regular convolution layer in UNet architecture^34^. We replace the regular convolution modules with DSC modules in our RSA model to reduce model parameter number and accelerate training speed.

Additionally, we use SA modules to further improve network performance. Both SA and Channel Attention (CA) modules are originally proposed as components of the Convolutional Block Attention Module (CBAM)^35^, a lightweighted attention method. As the training and testing input image sizes can be different, the CA module barely improves or even degrades the segmentation performance in our tests. We, therefore, only leverage the SA module in our model. The output of the SA module can be represented as *SA*(*x*) = *M*_*SA*_(*x*) ⨂*x*, where ⨂denotes element-wise multiplication, and *M*_*SA*_(*x*) is the 2D spatial attention map. To enable the element-wise multiplication, we broadcast the spatial attention map along the channel dimension to match the tensor size. The spatial attention values are determined by the average- and max-pooled features across channels. Specifically, the average- and max-pooled features are concatenated and convolved in a convolution layer:

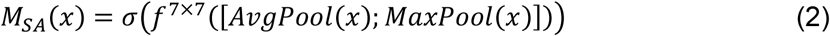

where σ(·) denotes the sigmoid function and *f* ^7×7^(·) denotes a convolution operation with a kernel size of 7×7.

We further use the residual connection to encapsulate the DSC and SA modules for direct information forward-feeding and back-propagation paths in our proposed deep network. Originally adopted to improve the image classification^36^, residual connection block has shown its promising efficacy for the biomedical image segmentation tasks^29,37^. Given the original network is denoted as *H*(*x*), its residual representation is *H*(*x*) + *x*. The residual connection in our proposed RSA block can improve the network performance without extra convolution layers.

(2)The second primary method development contribution is that we concatenate features from all decoders at different resolution levels as input to the output layer (i.e., orange arrows in Figure 1(C). In addition to features at the highest image level, the feature arrays in the lower image resolutions are leveraged in our model by convolving with a 3×3 filter for feature dimension reduction. The reduced features are resized to the highest image resolution by the bilinear interpolation before they are concatenated at the output layer. In contrast to FCN utilizing features from encoders^18^, our proposed model uses features from decoders. This design enables the output layer to make full use of multi-scale features and avoid the false negative problem with only a negligible increase in the parameter number. As there are multiple signal paths that lower resolution features from decoders can follow to reach the output layer in our model architecture, such a Multiple Up-sampling Path (MUP) mechanism is an effective solution to remedy the false negative problem observed in the UNet model in our study.

The architecture of our developed MUSA-UNet model is presented in Figure 1(C). Specifically, the MUSA-UNet consists of one input layer, four encoder-decoder pairs, and one output module. The encoders gradually decrease the image resolution by max-pooling layers while the decoders increase the image resolution by bilinear interpolation layers. In addition to the primary information encoding and decoding path, there are skip connections between the encoder output and the decoder input at each spatial resolution level. Therefore, there are two information sources provided to each decoder, one from a lower resolution decoder and another from the encoder output at the same resolution level. Note the feature representations from the lower resolution decoder are up-sampled and convolved before they are concatenated with the encoder output from the same resolution level. The outputs from distinct resolution levels are convolved and up-sampled before they are concatenated as the input to the output module.

### 2.3 Model implementation

Due to the overwhelming size of histopathology WSIs and the limited Graphical Processing Unit (GPU) memory size, deep learning models cannot be practically trained or tested on arbitrarily large images to achieve seamless segmentation. Therefore, we divide each WSI into image patches, apply trained models to individual patches, and assemble the patch-wise results.

A straight-forward partitioning strategy is to divide each WSI by a grid pattern. In that way, the segmentation output image can be produced by patch-wise segmentation results in the same spatial order of input image patches. However, the performance of this strategy could be degraded by the image patch border effect. We notice the prediction results of the same region in patches of varying sizes can be inconsistent, especially for those regions near patch borders. As deep learning analyses heavily depend on convolution operations and produce output patches of the same size as the input patches, padding methods for convolutions on pixels close to image borders are required^38^. The prediction results of pixels near patch borders are subject to the padded pixels and, therefore, can deviate from the ground truth.

To mitigate such image border effect, we have adopted a patch partitioning strategy that supports a seamless semantic segmentation^25^. Its overall schema is presented in Figure 2. First, we divide an input WSI by a regular grid pattern. To predict a target image patch in the grid, we extend its region scope before we provide it to our network MUSA-UNet for image segmentation. In Figure 2, the image regions denoted by dotted lines are the target image patches, while those in solid boundaries are extended counterparts. The margin for such an image patch expansion is set in such a way that prediction results of the original image patches are not influenced by padded pixels. After segmentation analysis by our deep learning network, we only retain the segmentation result of the interior regions associated with the original image patch region and assemble such results for the whole-slide segmentation maps by their spatial positions.

**Figure 2.**
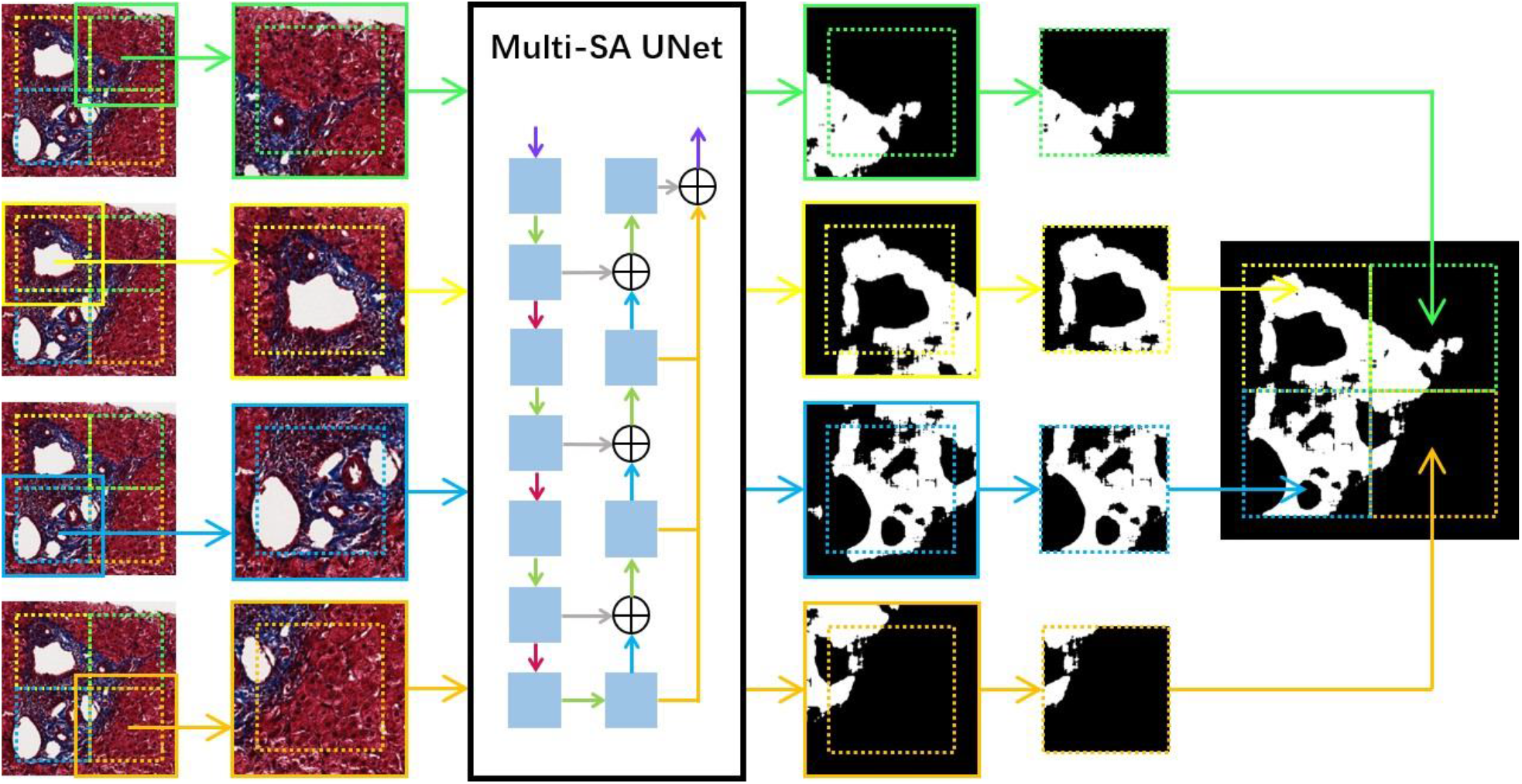
The patch partitioning strategy for seamless semantic segmentation in a large-scale image. To predict target patches in dotted lines, we extend these image patches before they are provided to our deep learning network for segmentation. The resulting segmentation output images are cropped back to the original patch size before the segmentation map aggregation.

In the testing stage, only image patches with enough foreground tissue (i.e., foreground patches) are expanded and provided to the trained network. Those with no significant tissue presence are skipped for the segmentation analysis, and the corresponding pixels in the resulting segmentation map are set to zero. For foreground patch recognition, we convert each image patch from the RGB to HSV color space and count the number of foreground pixels with a saturation value larger than 0.2. Those with more than 1% foreground pixels are considered as foreground patches. To accelerate the testing speed, we reduce the image resolution by 16 times before the foreground detection approach is applied in practice.

Note the strategy allows parallel computing on multiple GPUs. We implement codes in the Python 3.6 programming language and PyTorch 1.7.1 machine learning framework^39^ and run programs on two NVIDIA Tesla K80 GPUs. Balancing the tradeoff between network efficacy and computational efficiency, we design five image resolution levels in our model, with 64, 128, 256, 512, and 1024 filters from the highest to the lowest level, respectively. The loss function is the binary cross-entropy that can effectively reflect the pixel-wise difference between label and prediction. The model is trained with the Adam optimization algorithm^40^ for 40 epochs. The initial learning rate is set as 0.001 and the learning rate decay is 0.1 per ten epochs. In the testing stage, each image patch has 1,000 × 1,000 pixels, with an extended margin width of 140 pixels. Thus, each extended image patch has 1,280 ×1,280 pixels by size.

### 2.4 Portal tract guided fibrosis quantification

As reported in our prior study^1^, portal tract fibrotic percentage (i.e., portal tract fibrosis%) and average portal tract area derived from portal tract regions are correlated with Scheuer fibrosis staging. In this study, the Aperio ImageScope Positive Pixel Count (PPC) algorithm (Aperio Technologies Inc., Vista, CA) is applied to portal tract regions for quantification of the fibrous component in each portal tract by blue hue in the Masson’s Trichrome stain. After the fibrous components from the portal tract regions are measured by the PPC algorithm, the portal tract fibrosis% and the average portal tract area are computed. The portal tract fibrosis% is calculated as the proportion of the total fibrosis area in the total portal tract region area, while the average portal tract area is computed by dividing the total portal tract area by the portal tract region number in a slide. We further investigate the correlation of 1) Scheuer stage scores and average fibrosis areas; and 2) Scheuer stage scores and portal tract percentages (i.e., portal tract%), respectively. The average fibrosis area is computed by dividing the total fibrosis area by the portal tract number in a slide, while the portal tract% is the proportion of the total portal tract area in the total tissue area in a slide. We compute the total tissue area by subtracting the background pixel number from the total pixel number in an image.

### 2.5 Statistical analysis

In this study, statistical analyses are performed with Python3.6 and MATLAB R2021a (MathWorks Inc., Natick, MA). Precision, recall, F1 score, accuracy, Jaccard index, and Fowlkes–Mallows index are calculated. Correlations between variables are evaluated by linear regression and Spearman correlation. The student’s t-test is used to determine the statistical significance of the calculated Spearman correlation coefficients. A p-value less than significance level 0.05 is considered significant.

## 3. RESULTS

### 3.1 Dataset and annotations

This study includes 53 liver biopsies from patients who received liver transplantation. Annotations of the dataset are performed by two board-certified pathologists with GI/Liver pathology expertise (K.J. and A.B.F.). Biopsies are partitioned into training, validation, and testing dataset. Note all biopsies for the training and validation are mutually exclusive from those for the testing. Of all biopsies, 30 biopsies including 22 men and 8 women are used to generate image patches for model training and validation, with a mean ± standard deviation (S.D.) age of 54.5±6.9 years. We programmatically load manually annotated portal tract contours, calculate their bounding boxes, and divide them into patches of size 512×512 pixels. Additionally, we rotate image patches by 90, 180, and 270 degrees for training data augmentation. In total, we generate 6,012 image patches, with 80% and 20% for training and validation, respectively. The remaining 23 biopsies WSIs are allocated for testing, with 18 men and 5 women with a mean ± S.D. age of 51.8±7.7 years.

### 3.2 Portal tract segmentation results

We present in Figure 3(A) a typical portal tract region segmentation result by our proposed MUSA-UNet network. The model detected portal tract region borders are in yellow, while the ground truth portal tract regions are manually delineated and indicated by green borders in Figure 3(B). Such portal tract regions are automatically identified by binarization of the probability maps from our network in Figure 3(C). By visual assessments, we notice that the predicted region contours are highly concordant with the corresponding ground truth regions, suggesting the effectiveness of our proposed model. As detailed in the methods section, we divide each original WSI for testing into a set of patches and process them separately. Due to this partitioning step, portal tract regions close to image patch borders are subject to an image padding effect, resulting in inaccurate segmentation results. We present in Figure 4 portal tract segmentation results of two typical biopsy image regions divided with and without patch expansion partitioning strategy. We notice inaccurate segmentation results (by yellow arrows) when images are divided directly (blue dashed lines) Due to the border effect, portal tract regions on patch borders tend to be missed by the model. By contrast, the expanded patches by the patch expansion partitioning strategy are indicated by solid blue lines. This strategy substantially eliminates the segmentation errors by adding additional image margins to make the inception fields more informative and consistent.

**Figure 3.**
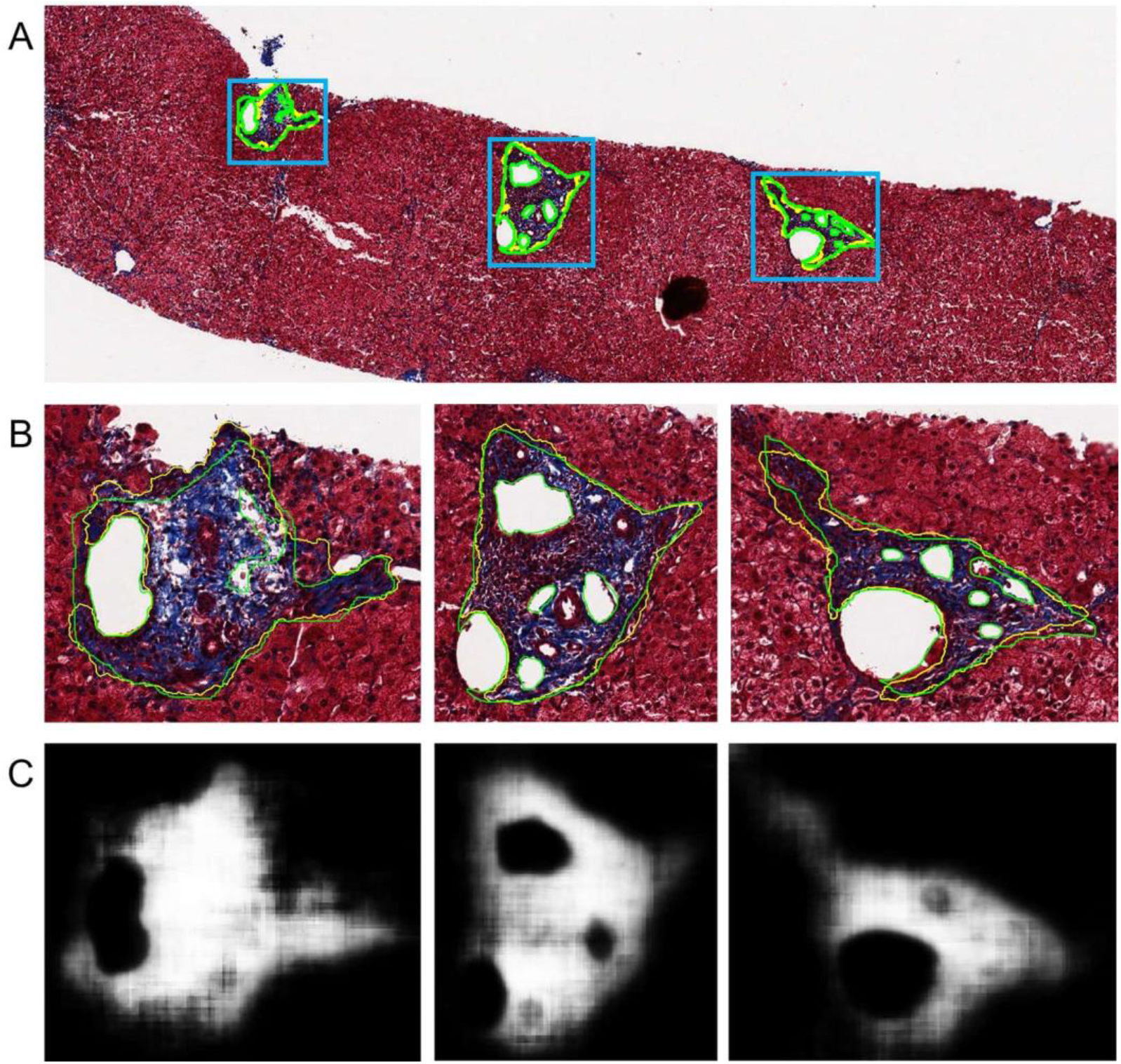
A typical portal tract segmentation result with a liver biopsy WSI. (A) Manual annotations (i.e., ground truth) and deep learning results of portal tract regions by the MUSA-UNet deep neural network are delineated in green and yellow, respectively; (B) Annotation and segmentation details are presented in close-up views; (C) The model generated prediction probability maps are presented for the same corresponding image regions.

**Figure 4.**
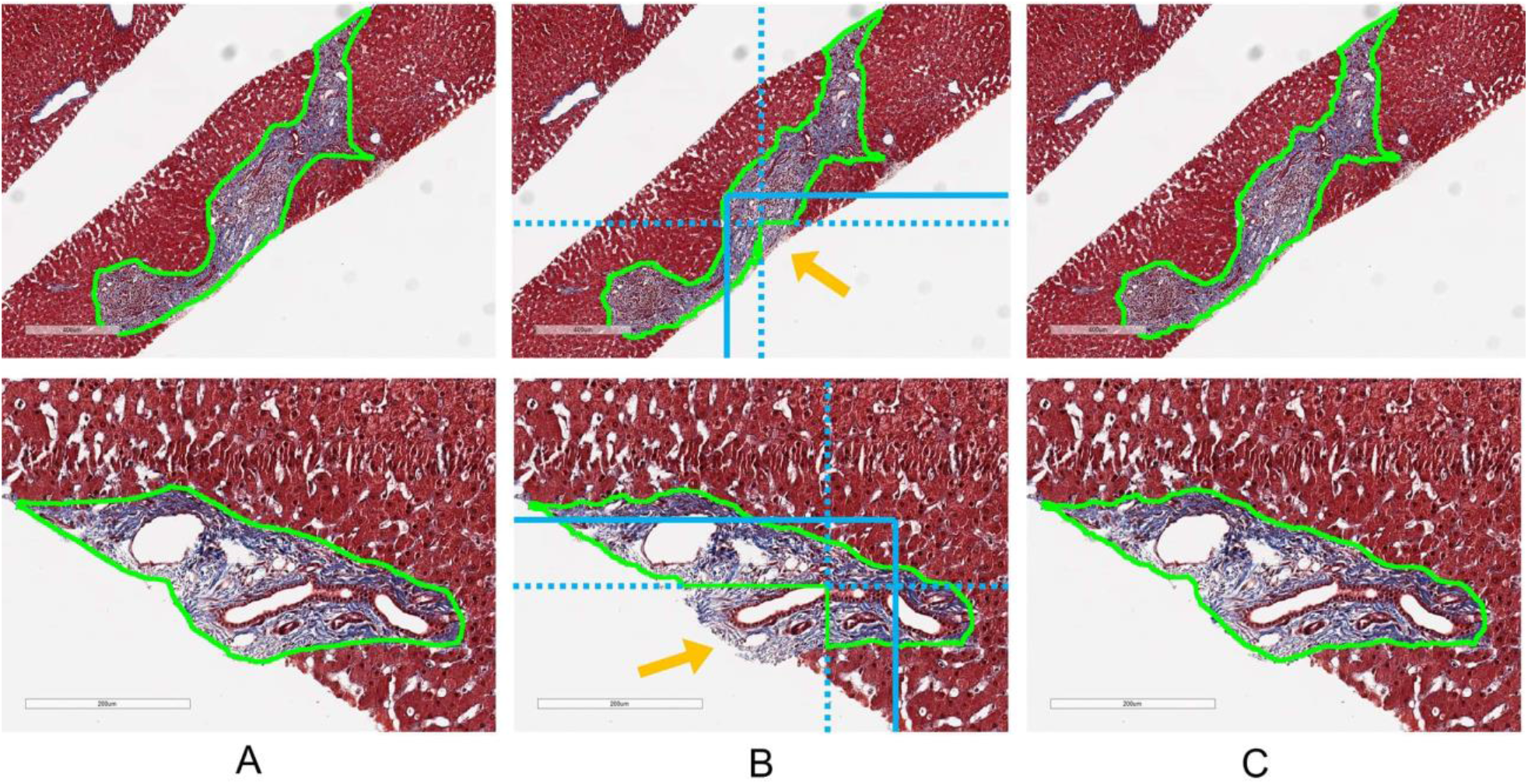
Comparison of portal tract segmentation results of two biopsy tissue regions with and without our partitioning strategy. (A) Ground truth portal tract contours are annotated by human experts; Portal tract segmentation results are presented when WSIs are simply divided into non-overlapping patches with their borders in blue dashed lines. The resulting segmentation defect is highlighted by a yellow arrow; (C) Portal tract segmentation results are demonstrated when the patch expansion partitioning strategy is used. The solid blue lines in (B) represent the borders of the expanded patches. With the expansion partitioning strategy, such negative border effects are successfully mitigated.

### 3.3 Deep learning model validation

In addition to qualitative assessments, we next validate our model quantitatively. We compare our proposed MUSA-UNet model with three widely used approaches, i.e., FCN^18^, UNet^25^, and DeepLab V3^41^. FCN and UNet have been widely applied to a large number of biomedical image segmentation tasks^9^. The DeepLab V3 model is derived from the FCN model, but with an atrous convolution^42^. This change expands the convolution perception field for enhanced segmentation accuracy without an increase in the parameter number. All the approaches are trained with the same training parameters and dataset as our model. We present and compare typical normal tissue segmentation results by these models in Figure 5. By visual comparisons, the segmentation results from our MUSA-UNet are more concordant with the ground truth than other methods. Additionally, we present and compare typical abnormal portal tract segmentation results in Figure 6. These abnormal portal tract types include portal tracts with (1) lymphoid aggregate, (2) ductular proliferation with minimal collagen, (3) edema, mild inflammation, and ductular proliferation, (4) features of acute cellular rejection, including mixed inflammatory infiltrate and ductitis, and (5) portal vein herniation and moderate chronic inflammation. Compared with FCN and DeepLab V3, our proposed MUSA-UNet demonstrates a better generalizability on abnormal portal tract segmentation.

**Figure 5.**
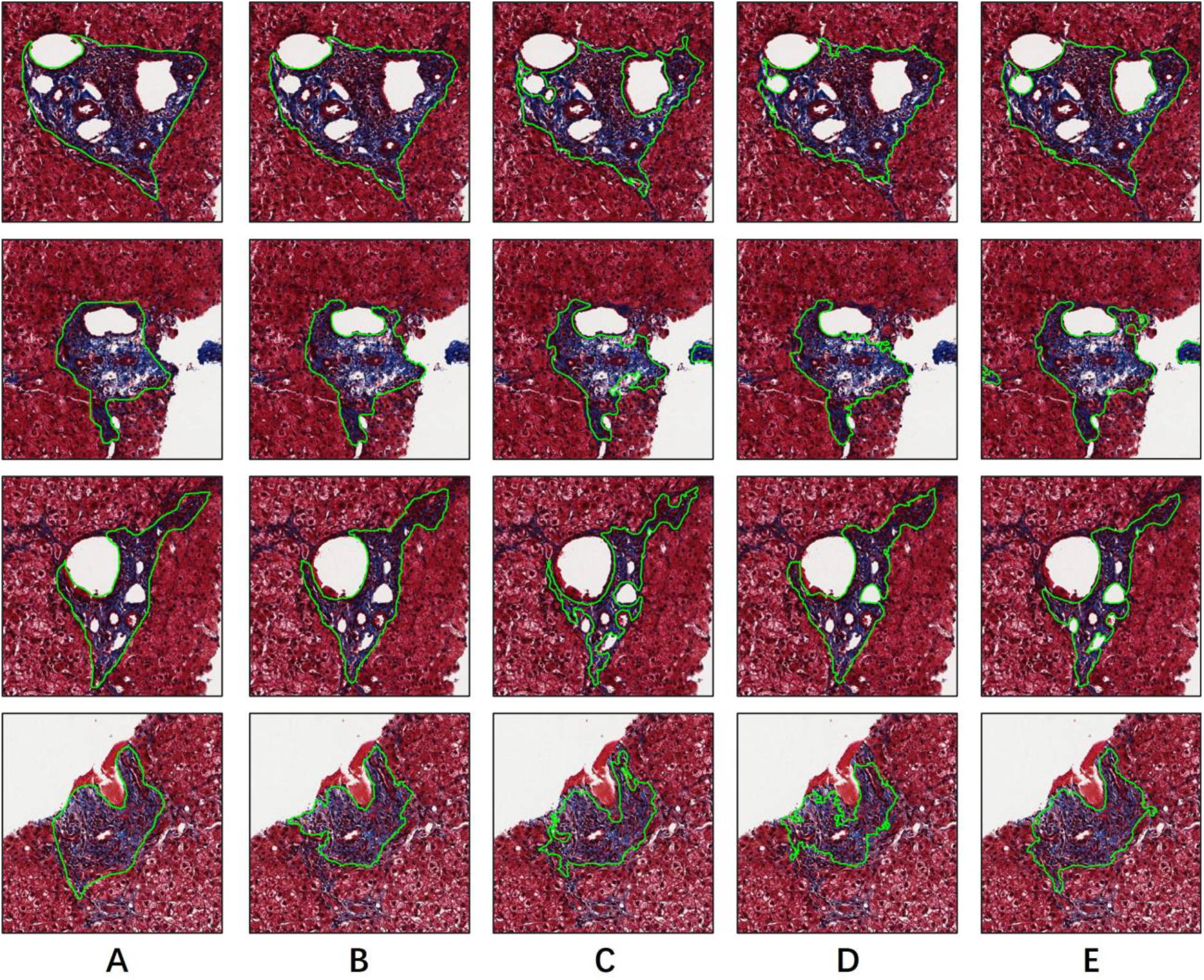
Qualitative comparison of deep learning models for normal liver portal tract segmentation. We present typical segmentation results of four normal liver tract regions by (A) human annotations (i.e., ground truth), (B) our proposed MUSA-UNet model, (C) DeepLab V3, (D) UNet, and (E) FCN, respectively.

**Figure 6.**
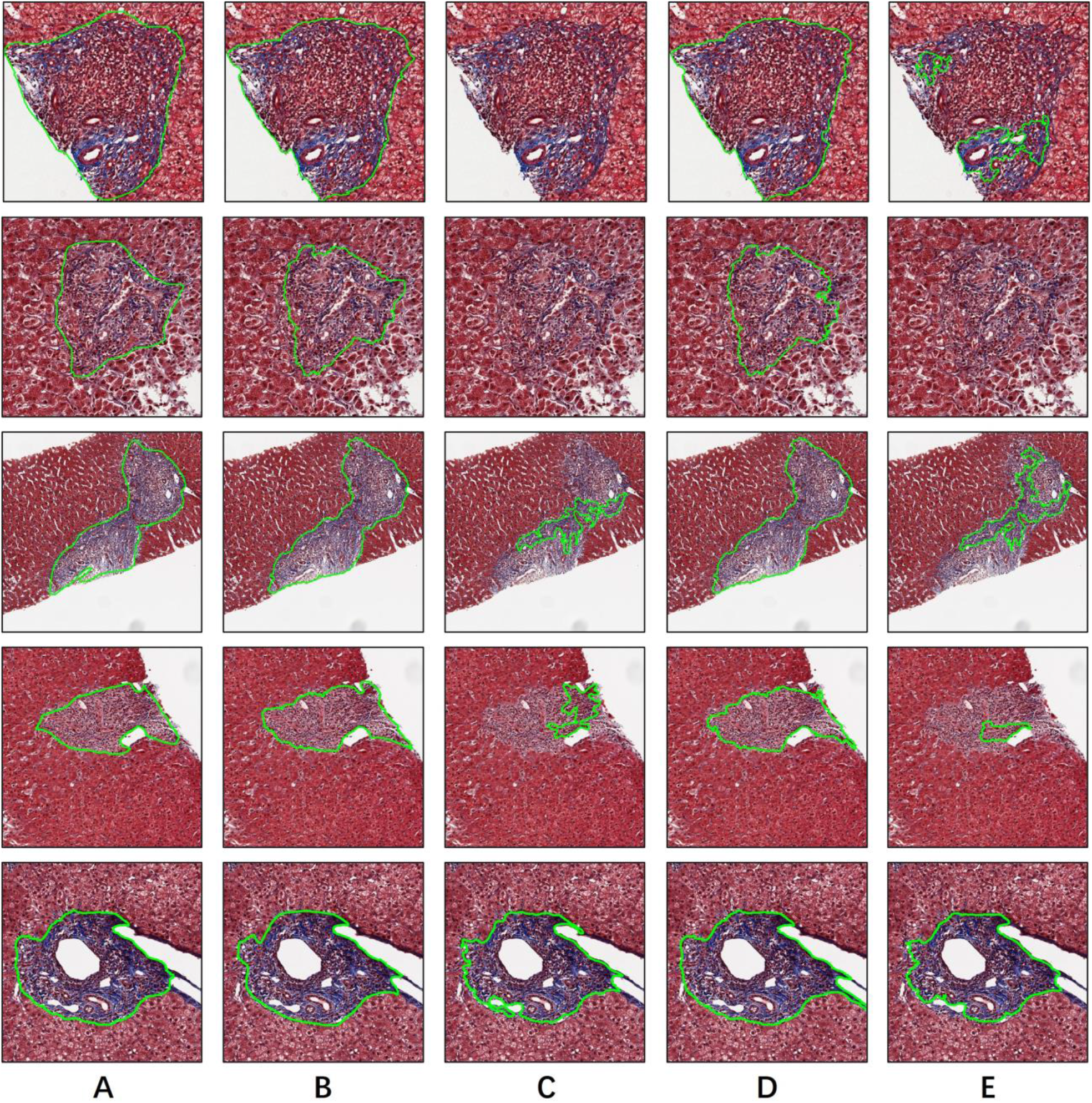
Qualitative comparison of deep learning models for abnormal liver portal tract segmentation. We present typical segmentation results of multiple abnormal liver tract regions by (A) human annotations (i.e., ground truth), (B) our proposed MUSA-UNet model, (C) DeepLab V3, (D) UNet, and (E) FCN, respectively. From top to bottom, we present abnormal portal tracts with (1) lymphoid aggregate, (2) ductular proliferation with minimal collagen, (3) edema, mild inflammation, and ductular proliferation, (4) features of acute cellular rejection, including mixed inflammatory infiltrate and ductitis, and (5) portal vein herniation and moderate chronic inflammation, lines).

Additionally, we compare segmentation results from different models with the ground truth from human annotations and quantitatively evaluate their performances. Compared to the ground truth, each pixel in the segmentation map is labeled as one of the four classes, True Positive (TP), False Positive (FP), False Negative (FN), and True Negative (TN). TP is the class for pixels that are correctly segmented as portal tract; FP is the label for pixels that are falsely recognized as portal tract; FN is the class for pixels that are missed as portal tract by mistake. Finally, TN is the label for pixels that are correctly recognized as non-portal tract. These four classes of pixels are illustrated in Figure 7. With these defined classes, we compute pixel-based evaluation metrics (each ranging from 0 to 1), including Precision (P), Recall (R), F1 score (F1), Accuracy (A), Jaccard index (JI), and Fowlkes–Mallows Index (FMI):

**Figure 7.**
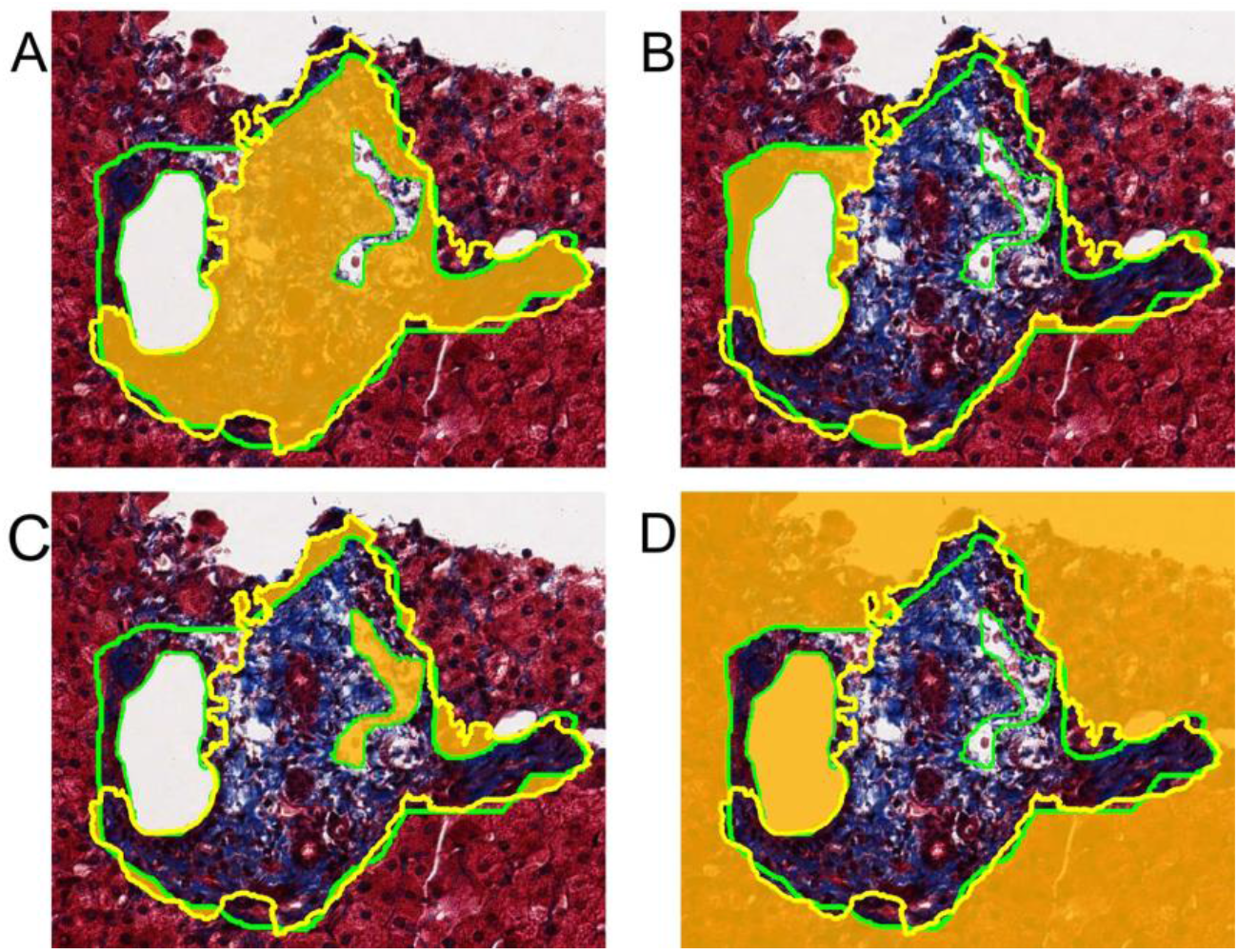
Pixel-wise segmentation labels for quantitative evaluation. Ground truth and deep learning segmentation results are represented by green and yellow contours. (A) TP is the class for pixels that are correctly segmented as portal tract; (B) FP is the label for pixels that are falsely recognized as portal tract; FN is class for pixels that are missed as portal tract by mistake; (D) TN is the label for pixels that are correctly recognized as non-portal tract.

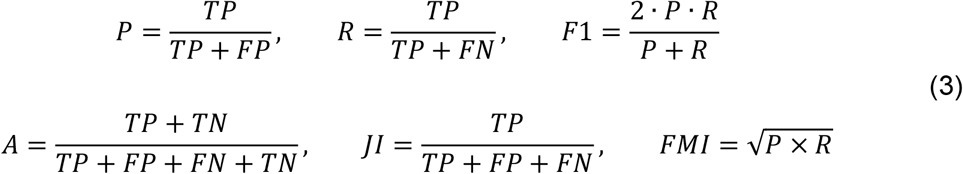

We present in Table 1 quantitative evaluation results of all models for comparison with Precision, Recall, F1 score, Accuracy, Jaccard index, and Fowlkes–Mallows Index. Although FCN has the best performance by Precision (0.9580), other methods (i.e., UNet, DeepLab V3, and MUSA-UNet) do not present significantly worse performances by paired sample t-tests with p-value 0.59, 0.30, and 0.29, respectively. By Recall, our MUSA-UNet demonstrates the best performance (0.8465) and a statistically significant performance difference compared with UNet, FCN, and DeepLab V3 with p-value 0.007, <0.001, and 0.03, respectively. By F1 score, our MUSA-UNet achieves the best performance (0.8857) and presents a statistically significant performance difference compared with UNet, FCN, and DeepLab V3 with p-value 0.01, <0.001, and 0.03, respectively. When assessed by Accuracy, our MUSA-UNet presents the best performance (0.8894) and a statistically significant performance difference compared with UNet, FCN, and DeepLab V3 with p-value 0.04, 0.002, and 0.04, respectively. By JI, MUSA-UNet has the best performance (0.8005) and presents a statistically significant performance difference compared with UNet, FCN, DeepLab V3 with p-value 0.01, <0.001, and 0.03, respectively. Our MUSA-UNet presents the best performance by FMI (0.9144) and a statistically significant performance difference compared with UNet, FCN, and DeepLab V3 with p-value 0.03, <0.001, and 0.05, respectively. We present the evaluation results in Figure 8(A) where evaluation performances of deep learning models for comparison are demonstrated by all six metrics. Note that MUSA-UNet presents fewer outliers than other methods, implying its strong stability. In Figure 8(B), we present and compare the Receiver Operating Characteristic (ROC) curves of MUSA-UNet, UNet, DeeplabV3, and FCN models, respectively. Of all these models, the proposed MUSA-UNet model achieves the largest Area Under the Curve (i.e., AUC=0.91).

**Table 1.**
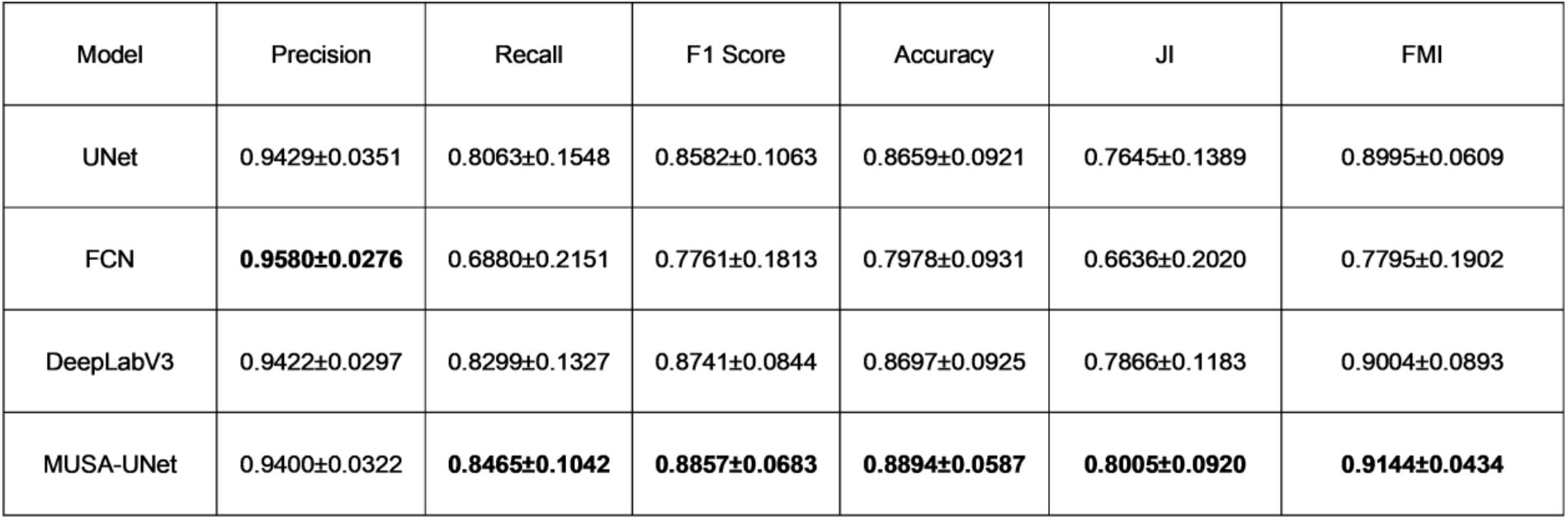
Quantitative performance comparison across the proposed MUSA-UNet model and other state-of-the-art segmentation models by multiple evaluation metrics (mean ± standard deviation).

**Figure 8.**
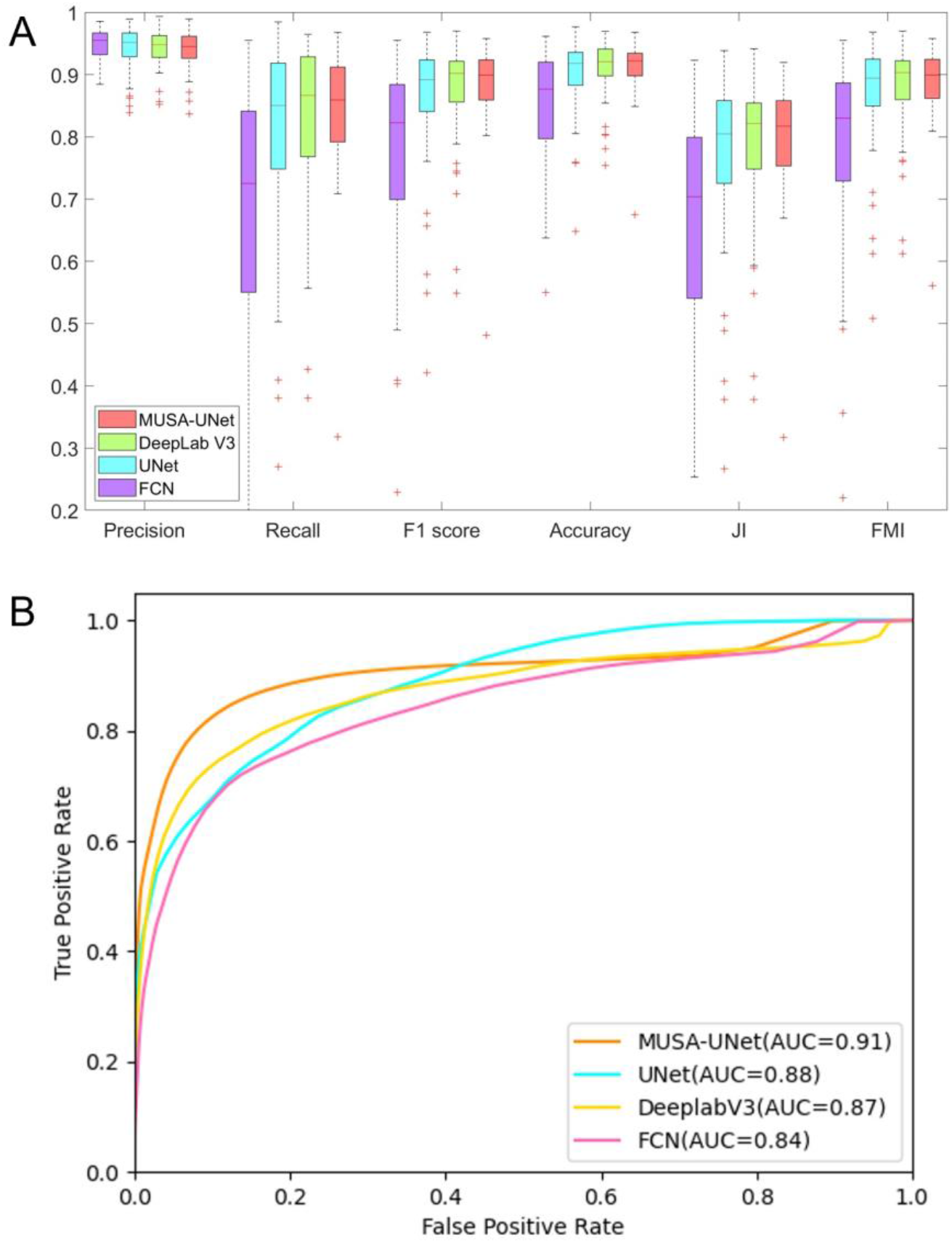
Quantitative comparison of deep learning models for liver portal tract segmentation. (A) Paired sample t-tests between the MUSA-UNet and other three widely used models (i.e., FCN, UNet, and DeepLab V3) suggest a statistically significant performance difference with p-values<0.05 by Recall, F1 score, Accuracy, JI, and FMI; (B) Of all deep learning models for comparison, our proposed MUSA-UNet achieves the best AUC with Receiver Operating Characteristic (ROC) curves.

### 3.4 Ablation study

To investigate the contribution of individual modules for portal tract segmentation, we carry out ablation experiments and present the ablation study results in Table 2. Small Attention UNet replaces convolution layers with two cascaded DSC modules and appends CBAM (CA+SA) blocks to DSCs^43^. Noticeably, model UNet+DSC+CBMA (i.e., SmaAt UNet) presents an inferior performance to that of the UNet model for the portal tract segmentation task. To identify the performance degradation reason, we remove either a SA or a CA module from the UNet+DSC+CBMA separately. The experimental results suggest that the DSC+CA dramatically decreases the performance while the DSC+SA improves Recall (0.8459), F1 score (0.8852), Accuracy (0.8804), JI (0.7992), and FMI (0.9107). We thus propose a new RSA block by retaining only one DSC module and encapsulating the DSC and SA models in a residual connection structure. In addition to an improved processing speed, the proposed RSA block achieves 0.8436, 0.8841, 0.8879, 0.7987, and 0.9136 by Recall, F1 score, Accuracy, JI, and FMI, respectively. The paired sample t-test between DSC+SA and RSA block results in a p-value less than 0.001, suggesting a comparable model performance of the proposed RSA block. To prove the effectiveness of the MUP mechanism, we add it to the original UNet and achieve improved performance by Recall (0.8157), F1 score (0.8666), Accuracy (0.8860), JI (0.7737) and FMI (0.8994).

**Table 2.**
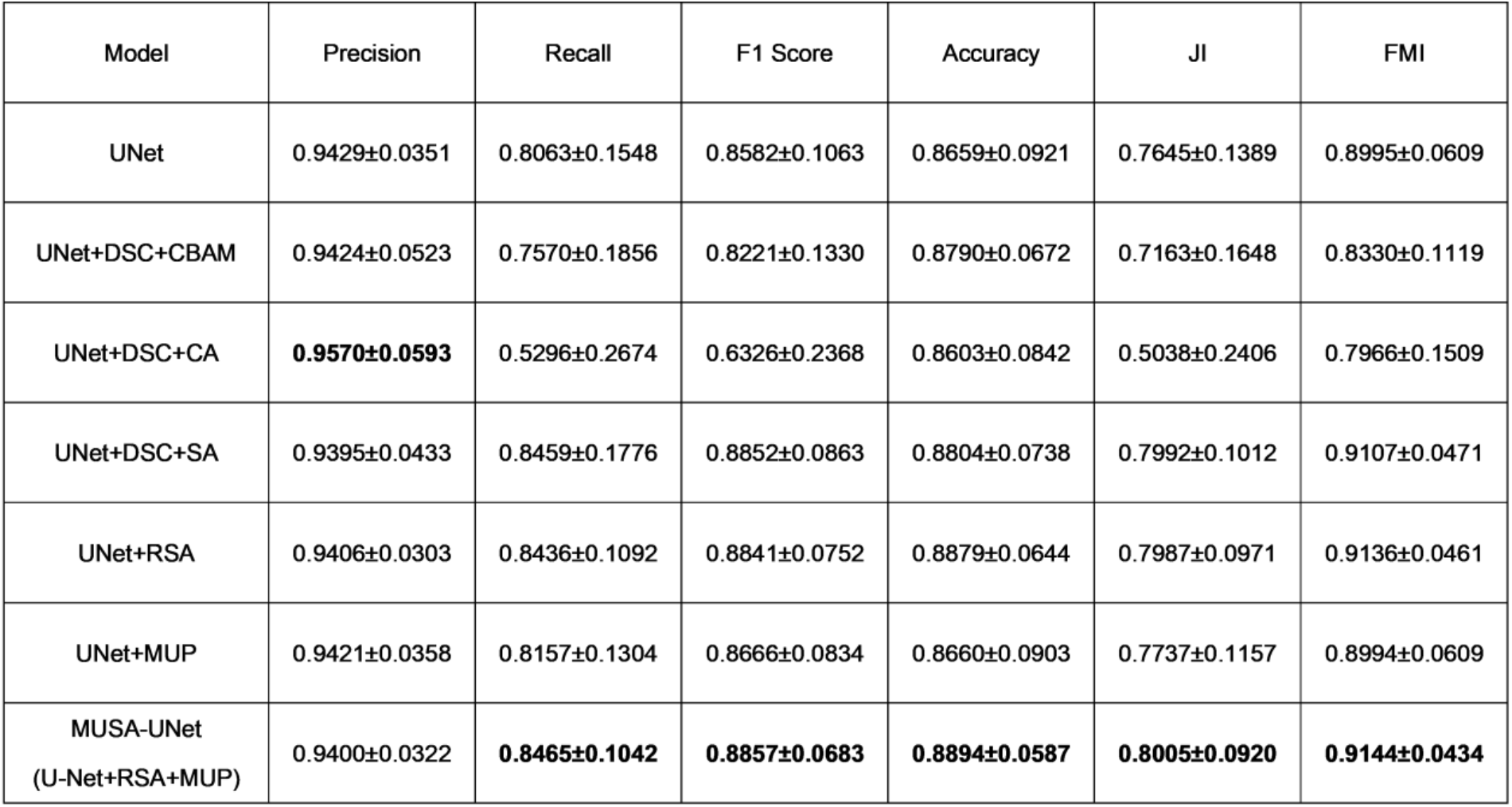
Quantitative model performance comparisons for the ablation study (mean ± standard deviation).

| Model | Precision | Recall | F1 Score | Accuracy | JI | FMI |
| --- | --- | --- | --- | --- | --- | --- |
| UNet | 0.9429±0.0351 | 0.8063±0.1548 | 0.8582±0.1063 | 0.8659±0.0921 | 0.7645±0.1389 | 0.8995±0.0609 |
| UNet+DSC+CBAM | 0.9424±0.0523 | 0.7570±0.1856 | 0.8221±0.1330 | 0.8790±0.0672 | 0.7163±0.1648 | 0.8330±0.1119 |
| UNet+DSC+CA | **0.9570±0.0593** | 0.5296±0.2674 | 0.6326±0.2368 | 0.8603±0.0842 | 0.5038±0.2406 | 0.7966±0.1509 |
| UNet+DSC+SA | 0.9395±0.0433 | 0.8459±0.1776 | 0.8852±0.0863 | 0.8804±0.0738 | 0.7992±0.1012 | 0.9107±0.0471 |
| UNet+RSA | 0.9406±0.0303 | 0.8436±0.1092 | 0.8841±0.0752 | 0.8879±0.0644 | 0.7987±0.0971 | 0.9136±0.0461 |
| UNet+MUP | 0.9421±0.0358 | 0.8157±0.1304 | 0.8666±0.0834 | 0.8660±0.0903 | 0.7737±0.1157 | 0.8994±0.0609 |
| MUSA-UNet<br>(U-Net+RSA+MUP) | 0.9400±0.0322 | **0.8465±0.1042** | **0.8857±0.0683** | **0.8894±0.0587** | **0.8005±0.0920** | **0.9144±0.0434** |

We present in Figure 9 typical segmentation results of four tissue regions by multiple ablated models for comparison. By visual comparisons, the segmentation results from DSC+CA are the worst as multiple portal tract regions are missing. This visual assessment conclusion agrees with the quantification analysis results. Although the result difference between the UNet+RSA and MUSA-UNet model is visually subtle, MUSA-UNet tends to produce smoother portal tract boundaries due to the new MUP design.

**Figure 9.**
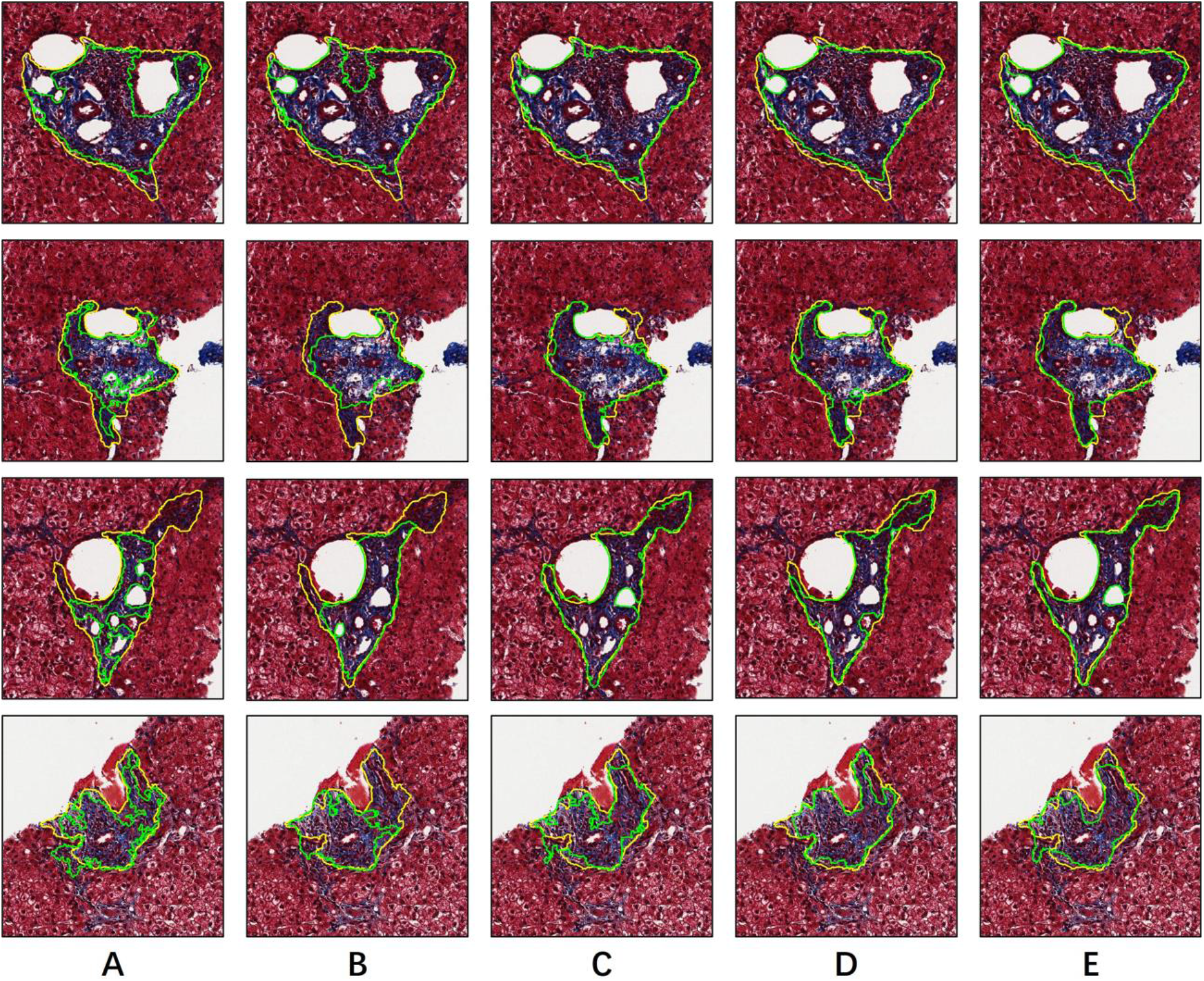
Qualitative comparison of ablated models for liver portal tract segmentation. We illustrate and visually compare typical tissue segmentation results of ablated UNet models (i.e., (A) UNet+DSC+CBAM, (B) UNet+DSC+CA, (C) UNet+DSC+SA, (D) UNet+RSA, and (E) UNet+MUP) in green and those of our proposed MUSA-UNet model in yellow.

### 3.5 Clinical correlation analysis

We investigate the correlation across measures of the portal tract area, the fibrosis area, and the clinical staging score. In addition to the ground truth established by the primary reviewers (reviewer 1: K.J., A.B.F.), a secondary board-certificated pathologist with GI/Liver pathology fellowship training (reviewer 2: N.S.) annotates portal tracts independently for this correlation analysis. Figure 10 demonstrates the multivariate analysis results with the linear regression and Spearman correlation. With manually delineated and MUSA-UNet predicted portal tract regions, we compute multiple measures, including portal tract fibrosis%, average portal tract area, average fibrosis area, and portal tract%. Additionally, we investigate and compare their correlations with the clinical Scheuer staging score (mean ± standard error: 0.85±0.23). The summary statistics for these measures are presented in Table 3. By Spearman correlation analysis, average portal tract area and portal tract fibrosis% derived from deep learning detected portal tract regions are correlated with clinical Scheuer staging score (R=0.681; p<0.001 and R = 0.335; p=0.02, respectively). When the MUSA-UNet derived measures are replaced with those from portal tract regions annotated by reviewer 1 and reviewer 2, average portal tract areas present comparable correlation relationships with clinical Scheuer staging score (i.e., R=0.680, p<0.001; and R = 0.574, p<0.001, respectively). With portal tract regions annotated by reviewer 1 and 2, the portal tract fibrosis% presents similar correlation relationships with clinical Scheuer staging score (i.e., R=0.437, p=0.002; and R = 0.326, p=0.016). Such comparable correlation results imply the good concordance between portal tract regions recognized by our proposed deep learning model and manual annotators. Figure 10 (A-B) demonstrates a strong correlation between human and deep learning derived measures, including portal tract fibrosis% and average portal tract area. Suggested by Figure 10 (C), the correlation between Scheuer staging score and average fibrosis area from deep learning identified portal tract regions is comparable to that between Scheuer staging score and average fibrosis area from human-annotated portal tract regions. By contrast, the correlation between Scheuer staging score and portal tract% from deep learning identified portal tract regions is stronger than that between Scheuer staging score and portal tract% from human-annotated portal tract regions in Figure 10 (D).

**Table 3.**
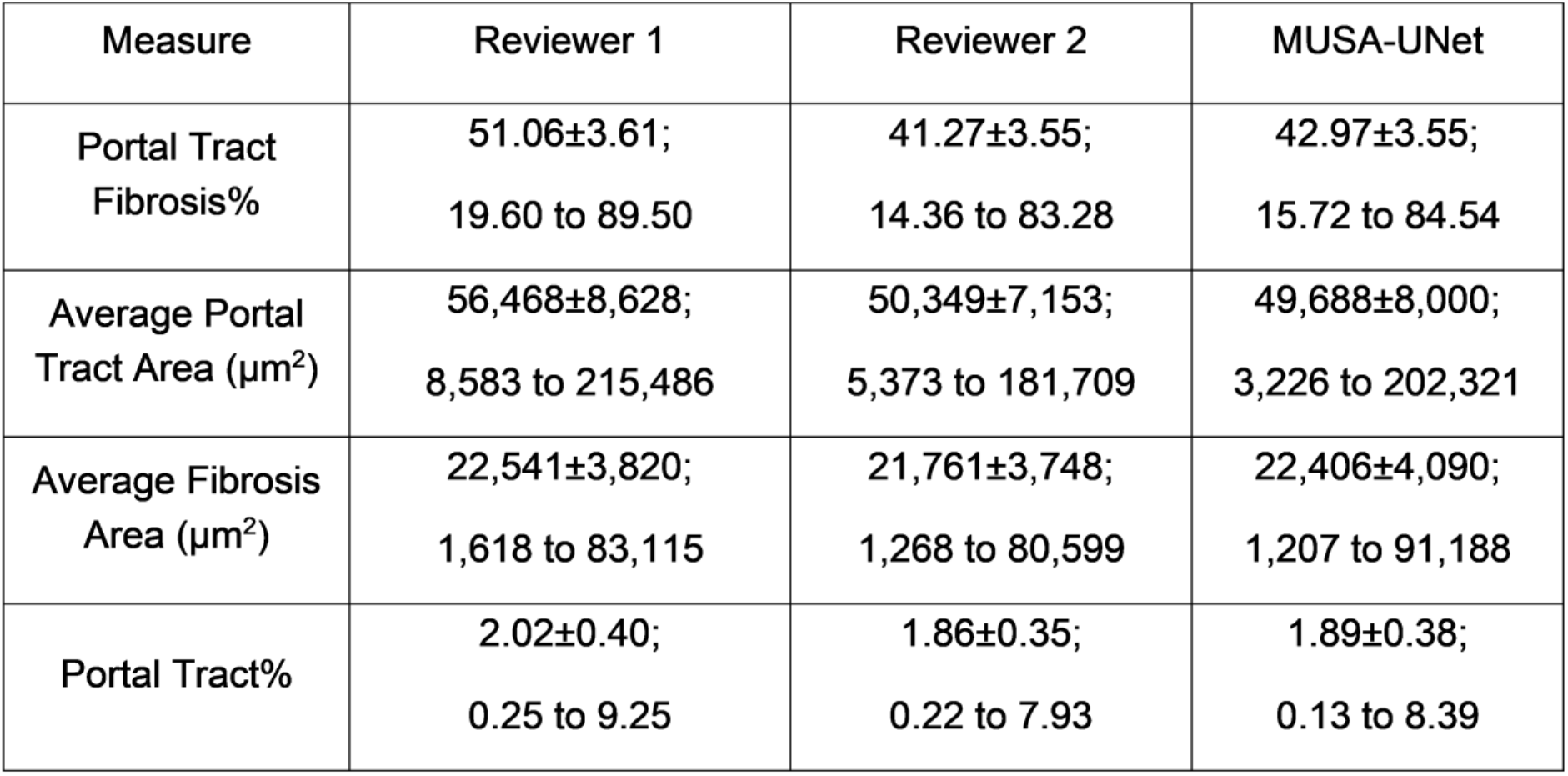
Summary statistics for multiple portal tract measures (Mean ± standard error; Range).

| Measure | Reviewer 1 | Reviewer 2 | MUSA-UNet |
| --- | --- | --- | --- |
| Portal Tract Fibrosis% | 51.06±3.61;<br>19.60 to 89.50 | 41.27±3.55;<br>14.36 to 83.28 | 42.97±3.55;<br>15.72 to 84.54 |
| Average Portal Tract Area ($\mu\text{m}^2$) | 56,468±8,628;<br>8,583 to 215,486 | 50,349±7,153;<br>5,373 to 181,709 | 49,688±8,000;<br>3,226 to 202,321 |
| Average Fibrosis Area ($\mu\text{m}^2$) | 22,541±3,820;<br>1,618 to 83,115 | 21,761±3,748;<br>1,268 to 80,599 | 22,406±4,090;<br>1,207 to 91,188 |
| Portal Tract% | 2.02±0.40;<br>0.25 to 9.25 | 1.86±0.35;<br>0.22 to 7.93 | 1.89±0.38;<br>0.13 to 8.39 |

**Figure 10.**
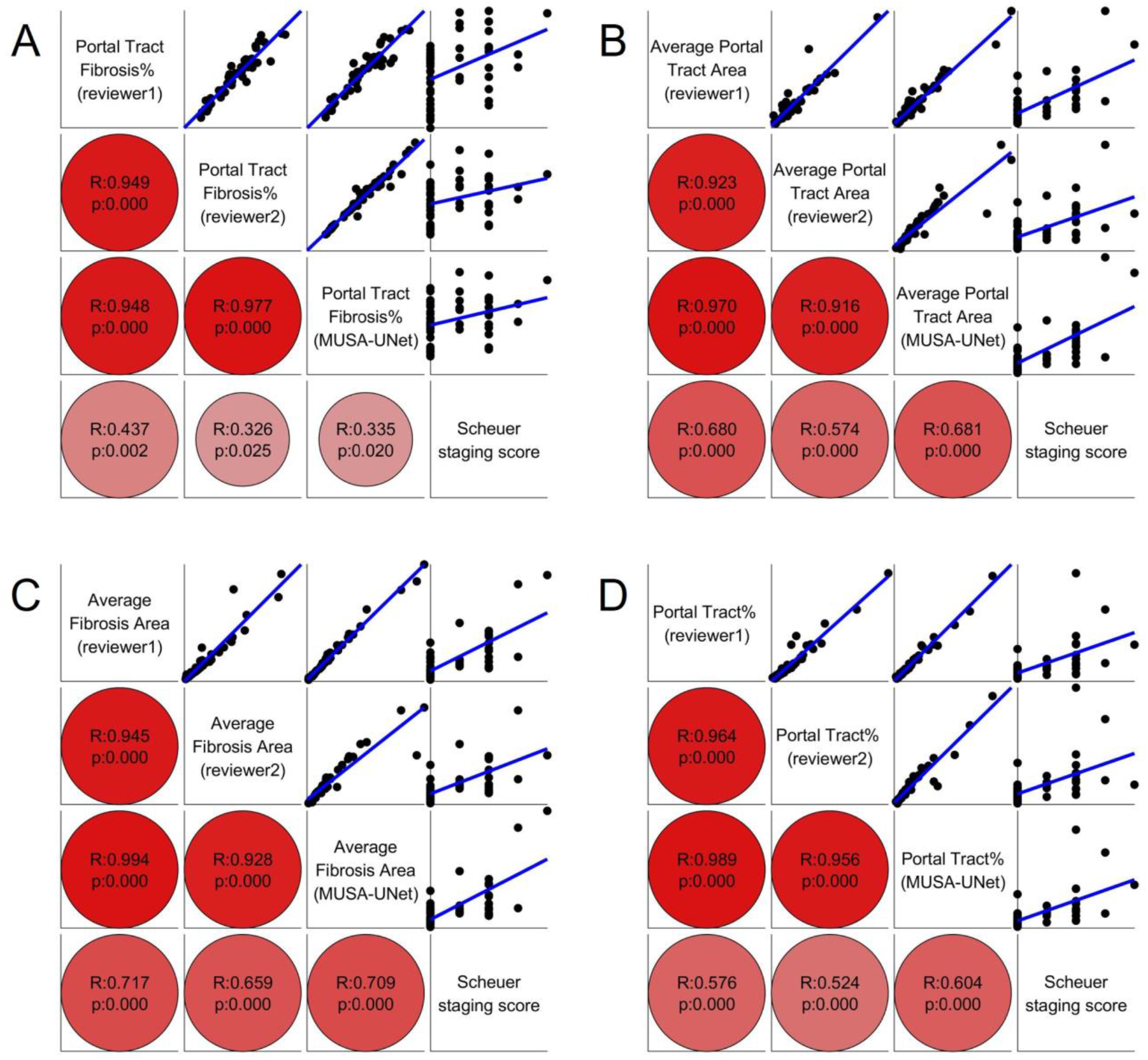
Multivariate correlation analysis across portal tract area, fibrosis area, and clinical staging score. The four subplots present the correlation analysis results between Scheuer staging score and (A) portal tract fibrotic percentage, (B) average portal tract area, (C) average fibrosis area, and (D) portal tract percentage, respectively. In each subplot, results from linear regression (top-right) and Spearman correlation (bottom-left) are presented to support the multivariate analysis. For Spearman correlation results, larger correlation coefficients and lower p-values are indicated by darker colors and larger circles.

Additionally, we demonstrate the differences in the clinical support between our proposed model and other methods for comparison. In Figure 11, we plot portal tract percentage populations by five Scheuer staging score groups, i.e., stage 0 to 4. Applied to the segmentation results from our proposed MUSA-UNet model, the analysis of variance (ANOVA) test suggests a significant difference in population means across staging groups with a p-value 1.44e-4. By contrast, p-values with results from UNet, FCN, and DeepLab V3 are 3.32e-4, 2.92e-2, and 7.70e-2, respectively.

**Figure 11.**
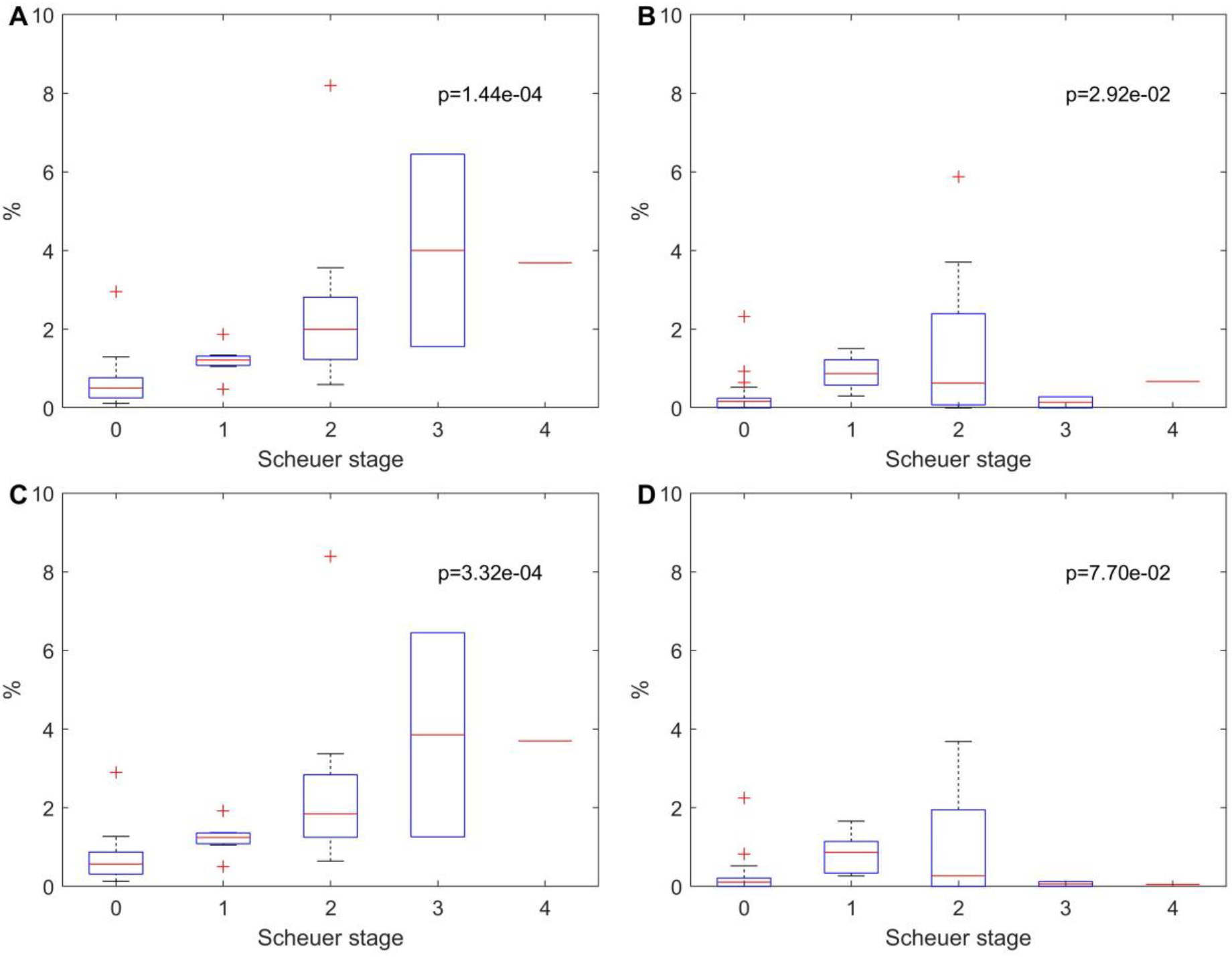
Comparison of deep learning models for clinical support. By ANOVA test, we present the significance of population mean difference across Scheuer staging groups with the portal tract percentage derived from segmentation results of (A) MUSA-UNet, (B) FCN, (C) UNet, (D) DeepLab V3, respectively.

## 4. DISCUSSION

### 4.1 Deep learning model optimization

Leveraging the UNet architecture as a building block, we develop the MUSA-UNet model for liver portal tract region segmentation with liver biopsy WSIs. To reduce the parameter number and accelerate the model processing speed, we replace the regular convolution layers in UNet with cascaded Depth-wise Separable Convolution (DSC) modules. By experiments, we notice Unet has a limited performance by Recall or JI. To further improve its performance, we include the attention mechanism in our model. Inspired by SmaAt UNet^43^, we first append a Convolutional Block Attention Module (CBAM) to the cascaded DSC modules (i.e., UNet+DSC+CBAM), leading to worse results. To investigate the cause of the model degradation, we change the original UNet by appending Channel Attention (CA) and Spatial Attention (SA) modules (i.e., two components in CBAM) to the cascaded DSC modules, respectively. The resulting UNet+DSC+CA model presents a degraded segmentation performance, while the UNet+DSC+SA model demonstrates an improved performance. Therefore, we only retain one DSC module, add a SA module, and encapsulate them by a residual connection block to make it more effective for back-propagation. This structure is defined as a Residual Spatial Attention (RSA) block. The resulting model (i.e., UNet+RSA) has fewer parameters, contributing to a better prediction performance and a faster execution speed.

The UNet architecture tends to focus on features derived from the highest image resolution level. By contrast, Fully Convolutional Networks (FCNs) only up-sample the output from the lowest image resolution layer (e.g., the FCN-32s model)^18^. Enlightened by these facts, we address the false-negative segmentation problem commonly seen around portal tract region boundaries by combining features from multiple image resolution levels for the probability map generation. Therefore, our network has a Multiple Up-sampling Path (MUP) mechanism, as there are multiple signal connections between lower resolution features from decoders and the output layer. By experimental results, the concatenated use of features from the top three image resolution levels significantly improves performance. Features from additional lower image resolution levels marginally improve the model performance, but at the cost of the increased model complexity.

### 4.2 Deep learning model complexity

In our model design, DSC modules are used to decrease the model parameter number. We present in Table 4 the parameter number and processing time cost of diverse models for performance comparisons. Compared with our proposed MUSA-UNet, the original UNet model has the same image resolution level number and the feature number in each level. The FCN model and the DeepLab V3 model are constructed on the base of the ResNet101 backbone^36^. By Table 4, the parameter numbers in the models without DSC modules (i.e., UNet, FCN, and DeepLab V3) are one order of magnitude larger than that of MUSA-UNet. This large difference in model parameter number has an important impact on the resulting processing speed. It takes about two hours for UNet to complete training with a data epoch on our current hardware setup, while the training time cost for the MUSA-UNet model is about 25 minutes. On average, it takes 23.4 ms for MUSA-UNet to predict a 512×512 image patch, promising to support an efficient segmentation analysis for clinical settings.

**Table 4.** Model comparison by parameter number and the average processing time cost for a 512×512 image patch. Processing time is based on the hardware setup described in our methods.

| Model | Parameter number | Processing time (ms) |
| --- | --- | --- |
| UNet | 37,384,833 | 30.5 |
| FCN | 51,938,881 | 31.1 |
| DeepLab V3 | 58,625,857 | 39.9 |
| MUSA-UNet | 9,123,958 | 23.4 |

### 4.3 Limitations and future work

Although our deep learning method is promising for automated segmentation of liver portal tracts in liver biopsy WSIs, there are some limitations to be addressed in the future. Our proposed RSA block only retains the spatial attention module from the CBAM. We specifically exclude the channel attention module as it tends to miss foreground when the foreground portion is limited in a given image patch. We hypothesize that these effects result from inappropriately chosen hyperparameters of the channel attention module (e.g., the hidden layer size). In addition, our current analysis on two-dimensional histopathology images of sampled tissue cuts is subject to the sampling bias, as portal tracts have three-dimensional morphology in liver tissues. In the future study, we will extend our analysis from two-to three-dimensional image space where portal tract structures can be captured in three-dimensional image volumes composed with WSIs of serial needle biopsy cuts. Finally, we plan to make our approach more generic to support a larger set of disease investigations using biomedical images.

## 5. CONCLUSION

Motivated by reducing intra- and inter-observer variability in the manual annotation process, we have developed an end-to-end deep-learning-based approach (MUSA-UNet) for portal tract region segmentation in liver whole-slide histopathology images. The developed deep learning model enables the production of seamless segmentation maps with accurate patch-wise prediction results. Built upon the UNet architecture, the proposed network significantly improves the portal tract segmentation efficiency and accuracy by the residual connection, the attention mechanism, depth-wise separable convolution modules, and the multiple up-sampling path mechanism. Our deep learning model is systematically validated and compared with widely used deep learning methods both qualitatively and quantitatively. Additionally, we present the effectiveness of our method by the correlation analysis with clinical staging scores. All experimental results demonstrate the efficacy of our proposed approach and suggest its promising clinical translational value for pathological review assistance.

## Data Availability Statement

Codes are available at Github repository: https://github.com/jkonglab/Liver_Portal_Tract_Segmentation

## Conflict of Interest Statement

The authors have declared no conflicts of interest.

## Acknowledgments

This research is supported in part by grants from National Institutes of Health 1U01CA242936, and National Science Foundation ACI 1443054 and IIS 1350885.

## Notes

### Competing Interest Statement

The authors have declared no competing interest.

